# Enhanced DNA Scanning and Target Specificity via Motif Sequestration

**DOI:** 10.64898/2026.10.12.758470

**Authors:** Jonathan S. Montgomery, Ehsan Akbari, Alexandar L. Hansen, Aparna Unnikrishnan, Michael G. Poirier, Mark P. Foster

## Abstract

Sequence-specific DNA-binding proteins must efficiently locate their target sequences within large, structurally heterogeneous genomes while avoiding nonproductive off-target interactions. Here, we show that an autoinhibitory conformational state of the site-specific recombinase Cre lowers nonspecific DNA-binding affinity to promote one-dimensional DNA scanning while suppressing off-target assembly of recombinogenic complexes. Recent studies indicate that the C-terminal segment of Cre transiently adopts a sequestered conformation that occludes the catalytic domain’s DNA-binding surface. Using NMR spectroscopy, single-molecule fluorescence microscopy, and optical trapping, we investigated how this conformational equilibrium regulates target search. NMR measurements reveal that the sequestered state predominates in the absence of DNA and persists at noncognate sites, whereas binding to cognate *loxP* sequences relieves autoinhibition. Single-molecule tracking demonstrates that full-length Cre undergoes processive one-dimensional diffusion and transitions to stable binding upon cognate-site recognition, whereas C-terminal truncation impairs scanning efficiency by arresting diffusion. These findings demonstrate how transient motif sequestration coordinates target search with site-specific complex assembly, suggesting a general mechanism by which DNA-modifying enzymes balance search efficiency and target specificity.

**Significance Statement:** Proteins that recognize specific DNA sequences must rapidly locate their targets within large genomes while avoiding nonproductive interactions with nonspecific DNA. How these competing demands are reconciled remains a fundamental question in molecular biology. We show that the recombinase Cre solves this problem through a reversible autoinhibitory mechanism in which a protein tethering motif is transiently sequestered within the DNA-binding interface. This sequestration promotes one-dimensional diffusion along DNA while suppressing inappropriate assembly at noncognate sites. Recognition of the correct target sequence releases the tethering motif, enabling formation of the active recombination complex. These findings reveal a general mechanism linking conformational dynamics, target search efficiency, and DNA-binding specificity.

## Introduction

Sequence-specific DNA-binding proteins are faced with the competing requirements of efficient target localization and stringent sequence discrimination within large and structurally heterogeneous genomes. Target localization is generally thought to involve mechanisms that combine three-dimensional diffusion with one-dimensional sliding or hopping along DNA (1–4). Yet, how sequence search functions to allow for rapid searching, limited off-target binding, and inappropriate functional assembly on non-cognate site, while having stable interactions upon localization with the cognate site, remains an open question. For enzymes whose activity depends on higher-order nucleoprotein complex formation, premature stabilization at non-cognate sites poses a particular challenge. We investigated the mechanism of target localization in the widely-used gene-editing enzyme, Cre recombinase.(5–8)

Originally evolved in bacteria to mediate circularization and replication of the P1 bacteriophage genome (7, 9), Cre recombinase has become an indispensable tool for precise DNA manipulation in mammalian systems (10). It enables recombination between sites separated by megabases (11) without producing cytotoxic double-strand DNA breaks (12, 13). As a member of the tyrosine family of site-specific DNA recombinases (YSSRs), Cre facilitates deletion, inversion, and insertion of genetic elements with high efficiency by assembling a tetrameric recombinogenic complex at its cognate *loxP* sites (**Figure 1A**)(9, 14, 5, 6). A widely used tool in biotechnology, Cre underpins tissue-specific and conditionally controlled gene knockouts (15, 16). More recently, engineered Cre variants are being explored as potential therapeutics, targeting DNA sequences of biomedical interest for the treatment of human diseases (17–20).

**Figure 1.**
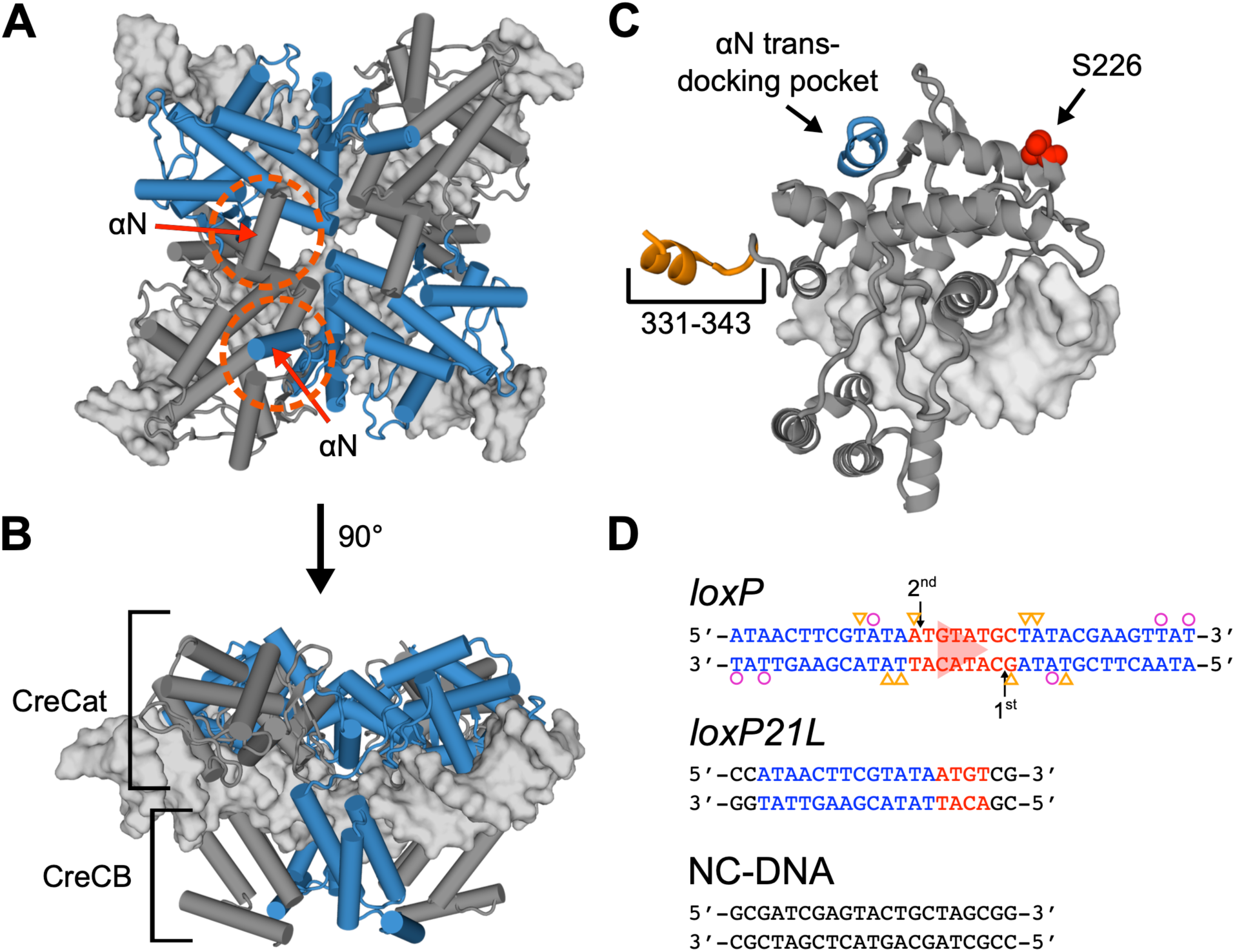
Architecture of Cre-*loxP* recombinogenic complexes. (A) Synaptic complex (PDB 2HOI) composed of two antiparallel *loxP* DNA duplexes (surfaces) and four Cre protomers (cartoons), with alternating elements adopting cleavage-competent conformations (blue). The C-terminal segment of each protomer forms helix αN, which docks in a clockwise fashion onto a neighboring protomer along the dimer interface or across the synaptic interface (red circles). (B) Side view illustrating how the N-terminal Core Binding (CreCB) and C-terminal Catalytic (CreCat) domains encircle the target DNA. (C) A Cre protomer and *loxP* half-site extracted from the synaptic complex. In the tetrameric complex, each protomer extends a C-terminal tethering motif (orange) and accepts one into its trans-docking site (blue), stabilizing the assembly. The site of a Ser226-Cys mutation introduced for this study is indicated. (D) The *loxP* DNA sequence features two 13-bp inverted recombinase binding elements (RBEs, blue) flanking an asymmetric 8-bp spacer sequence (red). Asymmetry in the spacer (indicated by the red triangle) results in the indicated strand cleavage order. Sparse base contacts in the major groove (triangles) and minor groove (circles) are indicated. A *loxP*21L half site contains an RBE (blue) and the left half of the spacer sequence (red). A noncognate DNA sequence (NC-DNA) used in this study with no homology with the cognate half site.

Understanding the determinants of site specificity in Cre recombinase is critical for its broader application, particularly in therapeutic contexts. Unlike nucleases whose specificity can be reprogrammed using oligonucleotide guides, the basis for site-selection by Cre remains poorly understood (6, 8, 21, 22). This gap persists despite extensive high-resolution structural analysis (5, 23, 24) and deep-sequencing studies (25–28).

Structural studies have provided limited insights into sequence determinants of Cre-*loxP* recognition. Cre comprises two structured domains, an N-terminal core binding domain (CreCB, residues 1-126) and a C-terminal catalytic domain (CreCat, residues 127-343), that bind DNA forming a C-shaped clamp (**Figure 1B**, **1C and S1**)(29). The catalytic domain contains the active site residues, including the eponymous tyrosine nucleophile, as well as a C-terminal helix-forming element αN, responsible for trans-docking in higher-order oligomeric species (**Figure 1A**, **1C**) (29–31). The *loxP* sequence comprises two 13-bp inverted repeat recombinase binding elements (RBEs) that flank an 8 bp asymmetric spacer (**Figure 1D**). Cre cooperatively dimerizes on these *loxP* sites and synapses in an antiparallel fashion with a second Cre_2_-*loxP* dimer to produce a tetrameric complex primed for recombination (29). The protein makes only eight major groove contacts and six minor groove contacts across the 34-bp *loxP* sequence, implicating a significant role for indirect readout in site recognition (23, 32). Library based high-throughput sequencing revealed selectivity for both contacted and non-contacted positions (26).

Solution nuclear magnetic resonance (NMR) and calorimetric experiments indicate that Cre exchanges between different conformations with distinct functional states that may influence DNA site selection (33, 34). Chemical shift and proximity data show that in the absence of DNA, the αN-forming C-terminal segment docks in *cis* over the DNA binding and active site (34). Removal of this C-terminal segment increases DNA-binding affinity, indicating that this cis-docked state autoinhibits interaction with both cognate and noncognate DNA substrates (33). Conversely, binding to *loxP* DNA correlates with the release of the C-terminus, allowing it to dock into neighboring protomers and promote the formation of higher-order oligomeric species (29, 34). These observations suggest that transient sequestration of the C-terminus plays the dual roles of preventing premature oligomeric assembly while regulating DNA binding and site selection by Cre.

In this work, we used single-molecule tracking and solution NMR to investigate whether sequestration of the C-terminal αN-forming segment of Cre recombinase functions to enhance the rate of translocation along DNA, facilitating the search for target *loxP* sites. Amide ^15^N NMR relaxation experiments in the absence of DNA reveal that the dominant *cis*-docked conformation of the C-terminus transiently samples an extended conformation. Single-molecule particle tracking experiments reveal that the C-terminus is required for efficient DNA scanning, while two modes of DNA scanning are detected upon encountering *loxP* sites. Together, these results connect microscopic internal motions to macroscopic DNA scanning efficiency and support a model in which domain sequestration facilitates efficient one-dimensional scanning, while coupled conformational changes signal target site recognition and set the stage for synapse formation and recombination.

## Results

### The Catalytic Domain of Cre Exhibits Two-State Exchange

Prior amide chemical shift and paramagnetic relaxation enhancement (PRE)-NMR data revealed spatial proximity between the C-terminus of Cre and the DNA binding surface of the catalytic domain, implicating the necessity of a dynamic process to adopt the conformations observed in crystallographic and cryo-EM structures (29, 34, 35). To characterize this process, we performed ^15^N chemical exchange saturation transfer (CEST) NMR experiments, which are sensitive to exchange dynamics on the millisecond timescale (36, 37), on the independently folding catalytic domain ([U-^15^N]-CreCat, residues 127-343) (34).

To generate CEST profiles, the intensities of signals in a ^1^H-^15^N heteronuclear two-dimensional correlation spectrum were monitored as a weak saturating radiofrequency field (B_1_) was scanned across ^15^N frequencies. For a subset of amide resonances, two dips were observed in the resulting CEST profile: a major dip at the B_1_ frequency matching the chemical shift of the observed signal (the major state), and a second smaller dip at an ^15^N frequency matching a minor state (**Figure 2A**). Signals mapping to the interdomain linker and helices αM and αN exhibit two dips in their CEST profiles, indicating conformational exchange (**Figure 2B**). This conformational exchange was quantified by global fitting CEST profiles to a two-state model using the Bloch-McConnel equations to extract chemical shifts, populations of the minor state, and the exchange rate (**Figure S2**) (37), yielding a minor state population *P*_B_ of 0.007 ± 0.001 and an exchange rate of 120 ± 20 s^-1^ (**Figure 2C**). With the major-minor equilibrium constant *K* = (1 − *P_B_*)/*P_B_*, the major state is favored by Δ*G* = −*k_B_T* ⋅ ln *K* ≈ 5*k_B_T*. These data indicate that the sequestered conformation of the Cre C-terminus is strongly favored in the absence of DNA, with transient sampling, on the millisecond timescale, of an alternate conformation, likely extended as observed in DNA-bound structures.

**Figure 2.**
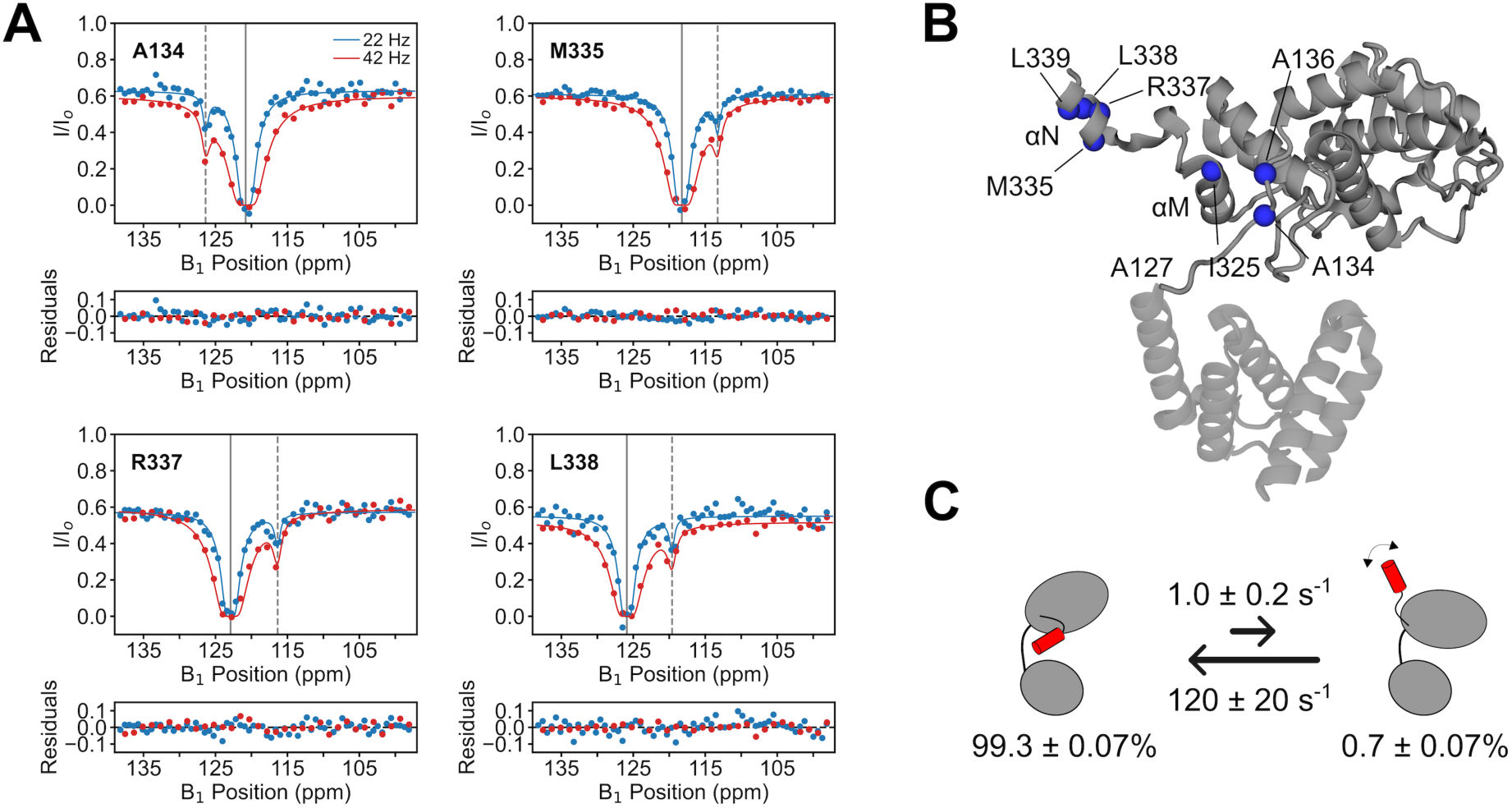
The sequestered C-terminal tethering motif of Cre is in a dynamic equilibrium with an extended conformation in the absence of DNA. (A) Representative backbone amide ^15^N NMR Chemical Exchange Saturation Transfer (CEST) intensity profiles obtained from spectra of the isolated catalytic domain (residues 127-343). Two-state exchange is indicated by the presence of a minor dip in the I/I0 profiles, with the minor state chemical shift indicated by a dashed line. An array of 2D ^1^H-^15^N correlated spectra were recorded by scanning ^15^N frequencies with 22 and 42 Hz saturating B1 fields. I/I0 ratios are relative to reference spectra recorded without an exchange time. Peak intensity profiles were fit by propagating the evolution of magnetization through a two-site exchange model. Below each CEST profile are residuals from best fits to the two-site exchange model. (B) Ribbon drawing of Cre recombinase as observed in synaptic complexes (chain A, 2HOI) with spheres highlighting residues that exhibit CEST profiles indicative of slow two-state exchange (panel A, and Figure S2). Helices αM and αN as indicated. (C) Schematic of the conformation exchange process probed in the CEST experiments with populations of sequestered and extended states, and exchange rates determined from global fitting. Forward and reverse rate constants were determined from fitted populations and exchange rates.

### Domain Sequestration Increases DNA Scanning Rates Along DNA

We hypothesized that domain sequestration would reduce off-target binding and thereby enhance target site search. To examine a role for this autoinhibited state in modulating one-dimensional diffusion of Cre on long DNA substrates, we employed a dual optical tweezers setup in combination with fluorescence microscopy (C-Trap, LUMICKS, Inc.) to track the motion of individual DNA-bound Cre molecules over time. To ensure single-fluorophore (Cy5) labelling of Cre molecules, a single cysteine variant was engineered by replacing all native cysteine residues (C49S, C56A, C155S, C240A) and reintroducing a single cysteine at a position (S226C, helix αI) predicted to be surface-exposed in free and DNA-bound states (**Figure 1C**). These mutations produced minimal structural or functional perturbations as assessed by two-dimensional ^1^H-^15^N NMR and *in vitro* recombination assays (**Figure S3**). To test the effect of autoinhibition on one-dimensional diffusion, we further engineered a single-cysteine derivative, Cre Δ330, which lacks the C-terminal helix (331–343) responsible for autoinhibition (33).

Using a microfluidics laminar flow cell to spatially segregate reagents during experiments, streptavidin-coated beads were trapped using optical tweezers (Figure S4). Single, double-stranded, end-biotinylated 48.5 kbp bacteriophage lambda DNA tethers were captured by the beads via biotin-streptavidin linkages. Single tethers were identified by fitting the force-extension curve to a worm-like chain model for a single DNA duplex (38). Captured single DNA tethers were positioned into the microfluidics channel containing 30 nM Cy5-labeled Cre for further study.

To monitor Cre interactions with DNA tethers, kymographs were collected with a time resolution (*dt*) ranging between 33-72 msec using a confocal beam scan along the contour length of the DNA tether held at a constant force of 10 pN, which has been shown not to significantly distort B-form DNA (38, 39). We observed frequent interactions of Cre with the tethered lambda DNA (**Figure 3A**).

**Figure 3.**
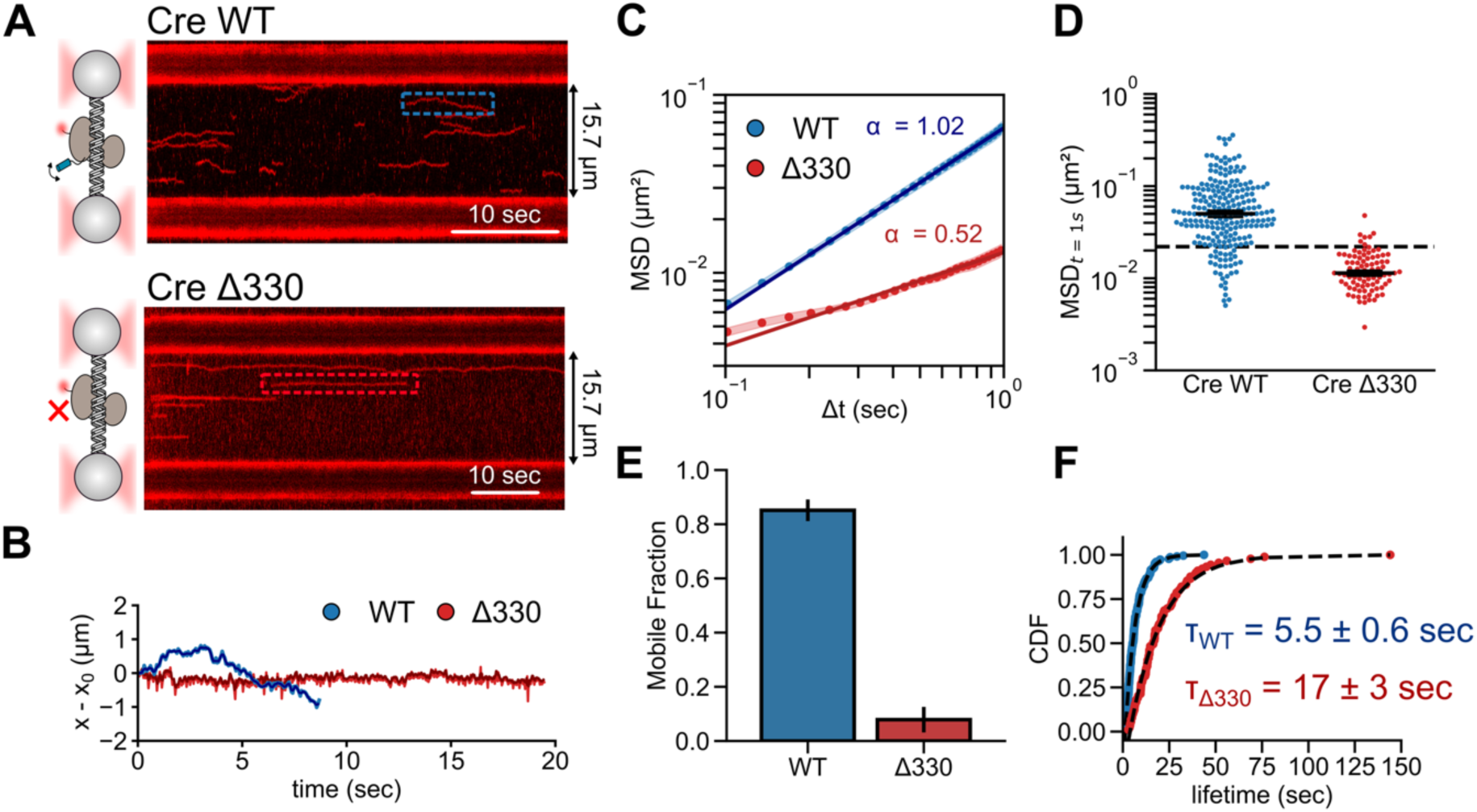
Motif sequestration enables DNA scanning by Cre. (A) Representative kymographs tracking Cre WT (top) and Cre Δ330 (bottom) particles on DNA; Cre Δ330 lacks the C-terminal *trans*-docking motif. (B) Example traces showing displacement from initial binding positions for WT (blue) and Δ330 Cre (red). (C) Mean squared displacement versus lag time Δ*t* = *n* ⋅ *dt* was computed for all traces with a time resolution of 33 ms; linear fits yielded values of *D* and *α* of 0.034 and 1.02 for WT Cre, and 0.0066 and 0.52 for Δ330. *D* is the diffusion constant with units of µm^2^ s^α^. (D) Swarm plot of the MSD at a time lag of 1 second for each trace (N_WT_ = 222, N_Δ330_ = 91); error bars represent the median ± standard error of the mean for each construct. Dashed line indicates cutoff for particles considered mobile. (E) Percent of particles classified as mobile based on minimal cutoffs of MSD_t=1sec_ = 0.11 and α = 0.2 (see text). (F) Cumulative distribution plots (CDF) of bound lifetimes τ of all traces, fit with a single exponential decay function, *A*(*t*) = 1 − exp(−*τ*/*t*) (dashed lines); uncertainties were obtained by bootstrapping.

Visual comparison of kymographs generated from Cre WT and Cre Δ330 reveals that autoinhibition significantly impacts its mobility on DNA (**Figure 3A**). The WT protein is observed to diffuse away from the origin of binding on the timescale of seconds. In contrast, Cre Δ330 molecules appear immobile over a similar time frame and remain colocalized at the initial encounter position.

To quantify diffusion of Cre on DNA, we tracked the position of each Cy5-labeled Cre in each kymograph over time (e.g., **Figure 3B**). The mean squared displacement (MSD) for all traces was then computed as a function of lag time Δ*T* = *n* ⋅ *dt* (Figure 3C), using equation 1. Here, *dt* is the sampling delay, *N* is the total number of positions in the trace, and *i* denotes the discrete position in the trace:

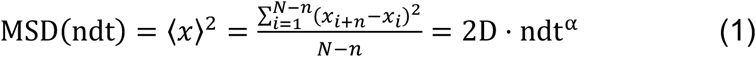

For one-dimensional diffusion, the MSD is proportional to the diffusion coefficient (*D*), while a power-law exponent that differs from unity (*α ≠* 1) accounts for deviations from random walk behavior. Values less than 1 approximate constrained diffusion, whereas values greater than 1 indicate energy-coupled translocation (40). In principle, *D* and *α* can be obtained from the power relationship between MSD and Δ*T* (**Figure 3C**), although these estimates may be distorted by positional uncertainty and finite camera exposure (41).

This analysis yielded an *α* = 1.02 ± 0.01 for WT Cre, consistent with random walk behavior, whereas the data for Cre Δ330 resulted in an apparent *α* = 0.52 ± 0.01, consistent with limited mobility and a dominant contribution from positional uncertainty. Evaluation of the apparent *D* at Δ*T* = 1 s implies a value of 34 ± 2 x10^-3^ μm^2^ s^-1^ for WT Cre (**Figure 3D**). For Cre Δ330, given the low α, the value at 1 s likely reflects fluctuations of tether, rather than sliding along it. Applying minimum cutoffs of 0.011 µm^2^ s^-1^ for MSD and 0.2 for *α* to classify diffusive particles (42), we find that 85 ± 4 % of WT Cre traces can be categorized as mobile over the sampled timescale, compared to only 8 ± 5 % for Cre Δ330 (**Figure S5, Figure 3E**). These data indicate that WT Cre undergoes random walk diffusion on DNA, whereas deletion of the C-terminus renders the protein effectively immobile.

Domain sequestration additionally influences the residence time of Cre on the lambda DNA tether. The lifetimes for individual traces were plotted as cumulative distribution functions and fit with a single exponential decay model, *CDF*(*t*) = *CDF*_0_(1 − *e*^−*t*/*τ*^, where τ is the characteristic dwell time. The fitted values were 5.5 ± 0.6 sec and 17 ± 3 sec for WT Cre and Cre Δ330, respectively (**Figure 3F**). The shorter dwell time for the diffusing full-length protein is consistent with its weaker DNA binding affinity compared to the immobile Δ300 deletion construct (33). Kymographs collected at increasing excitation power indicate that the observed characteristic lifetimes are not limited by dye photobleaching (Figure S6). Based on the fitted diffusive parameters, the mean linear distance scanned per binding event can be determined as (40):

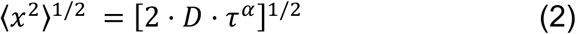

For the WT protein, this analysis indicates a typical scanning distance of ∼600 nm (1800 bp) per the typical bound lifetime of 5.5 s, whereas Cre Δ330 is essentially immobile over a similar timeframe.

### Domain Sequestration Enables a Unique Scanning Conformation on Noncognate DNA

To gain structural insights into the role of the C-terminal sequence of Cre in efficient DNA scanning, we recorded two-dimensional ^1^H-^15^N TROSY-HSQC spectra of uniformly ^15^N-labeled WT and Δ330 Cre alone and in complex with noncognate DNA (NC-DNA) (**Figure 4A**). Under conditions of the NMR experiments, both proteins are fully bound, as reflected in the broadened line shapes and insensitivity to excess DNA (**Figure 4A**).

**Figure 4.**
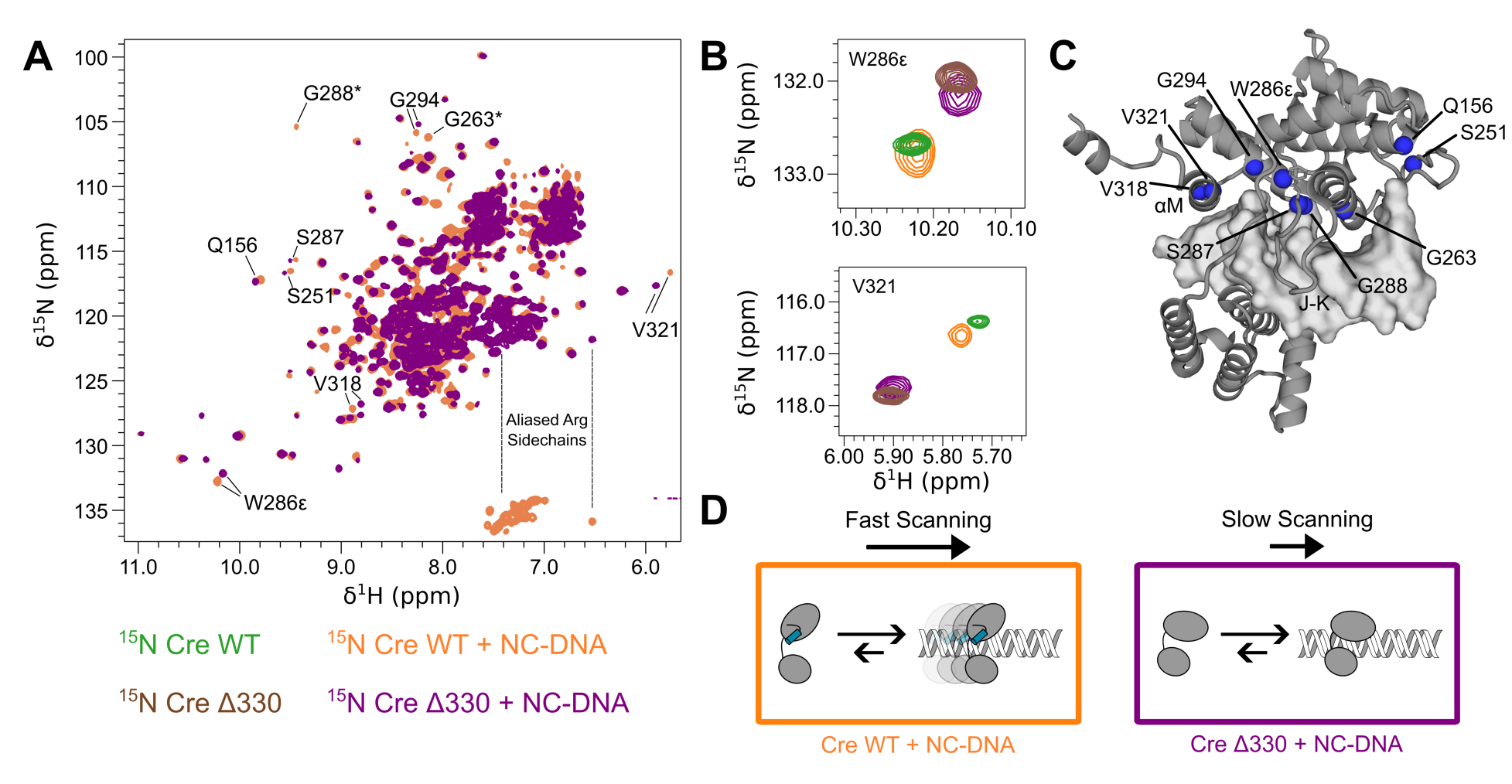
Motif sequestration enables a unique scanning conformation on noncognate DNA. (A) Overlay of ^1^H-^15^N TROSY-HSQC spectra of WT Cre (orange) and Cre Δ330 (purple) bound to noncognate DNA, with isolated resonances labeled; asterisks note signals for which the corresponding resonance is missing in the Δ330 spectrum. Extensive spectral differences indicate structural differences in the protein-DNA complexes. (B) Spectral overlays of apo WT Cre (green), WT Cre bound to noncognate DNA (NC-DNA, orange), apo Cre Δ330 (brown), and Cre Δ330 bound to noncognate DNA (purple). Similarity of WT Cre in the absence and presence of non-cognate DNA, and of Δ330 in the absence and presence of DNA indicates that the sequestered conformation persists in DNA-bound WT Cre (C) Positions of residues associated with motif sequestration mapped onto the structure of Cre bound to DNA (chain A, 2HOI; blue spheres). (D) Schematic summarizing a structural interpretation of the NMR spectra: WT Cre remains in a sequestered conformation on noncognate DNA, whereas Cre Δ330 adopts a conformation in which both domains more fully engage with DNA.

Full-length and Δ330 Cre interact differently with non-cognate DNA. NMR spectra of the Cre catalytic domain, and its Δ300 deletion bound to a cognate DNA half-site are nearly identical, except for additional signals corresponding to the C-terminal sequence deleted in Δ330 (**Figure S6**). In contrast, spectra of full-length Cre and Δ330 bound to non-cognate DNA are highly dissimilar, indicating that the C-terminus influences the conformation adopted when bound to noncognate DNA. Although complete backbone resonance assignments are not available for DNA-bound Cre, many resolved resonances that shift or disappear in the Cre Δ330 spectrum can be assigned to residues proximal to the autoinhibition docking/DNA binding interface (**Figure 4A** and **C**). These changes suggest that structural differences extend throughout the catalytic domain rather than being localized near the deletion site, implying that the *cis*-docked conformation may persist on non-cognate DNA substrates.

Congruence between NMR spectra of Cre alone and bound to non-cognate DNA supports a role for domain sequestration of the C-terminal segment. We focused our analysis on well-resolved backbone amide resonances from residues in the αM helix and flanking the J-K loop that are sensitive to the conformation of the C-terminal segment (**Figure 4B**)(34). For the representative resonances highlighted, we observe that signals for WT Cre overlay well in the absence and presence of non-cognate DNA (green, orange). This congruence implies that Cre does not fully engage with non-cognate DNA sequences and remains in a conformation in which the C-terminus remains docked *in cis*. However, signals from the same residues in Cre Δ330 bound to NC-DNA appear displaced, reflective of a more intimate DNA recognition state. These observations support a model in which domain sequestration enables DNA scanning by adopting a scanning-competent state on noncognate DNA (**Figure 4D**).

### Cognate site recognition arrests diffusion

To determine how encounters of Cre with its cognate *loxP* sequence influence sequence scanning, we designed a DNA tether containing a single *loxP* half site (**Figures 5A** and **S7**). Tethers with cognate and non-cognate inserts were held at a constant tension of 10 pN and kymographs were recorded. Although the spatial resolution of these experiments is insufficient to determine binding to the half-site from proximity, for the tethers containing the cognate *loxP* half site, Cre molecules were observed to localize at the center of the DNA tether with a range of lifetimes that were significant longer than tethers lacking the cognate site (**Figure 5B-D**). In fact, Cre particles were observed to bypass the center point of the tethers with dwell times less than one second with the non-cognate site (**Figure S8**). Figures 5 B and C illustrate traces with localization lifetimes of ∼5 and ∼65 seconds, respectively, while Figure 5 D illustrates an example in which Cre appears to only transiently associate near the *loxP* half-site.

**Figure 5.**
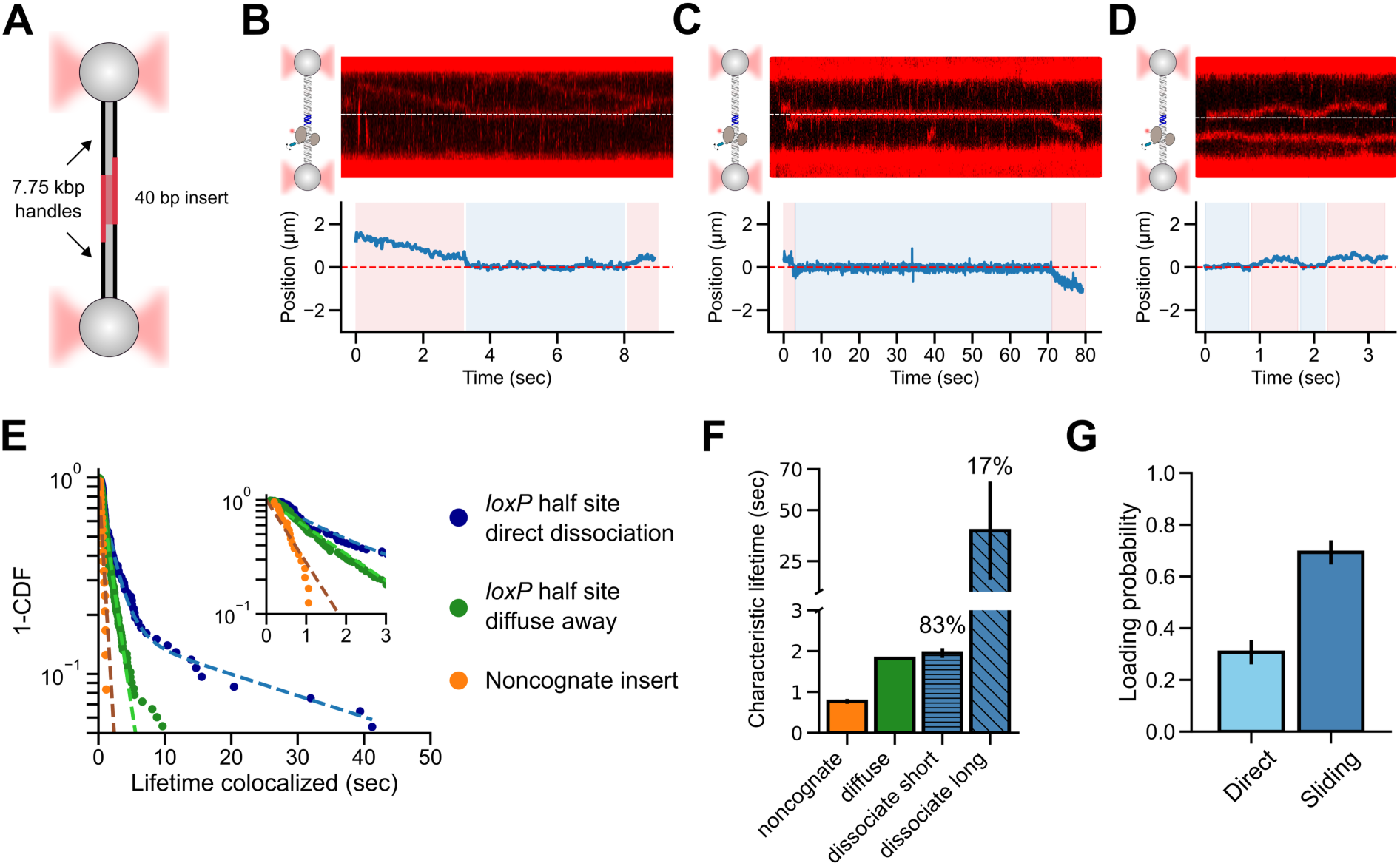
Cre diffusion is arrested at cognate DNA sites. (A) Designed DNA tethers containing inserts with either a cognate *loxP* half site or noncognate DNA. (B-D, top) Representative kymographs of Cre diffusing on a DNA tether containing a cognate *loxP* half-site sequence. (B-D, bottom) Tracked particles from corresponding kymographs (blue) plotted against the position of the *loxP* half-site (red dashed line). Time spans in which the particle is co-localized with the *loxP* half site are highlighted blue. (E) Cumulative distribution function (CDF) plots of the lifetimes of particles colocalized at the *loxP* half site (blue, direct dissociation, N = 177; green, linear diffusion, N = 82) or the equivalent position on tethers with noncognate DNA (orange, N = 23); inset is an expanded view of the data. (F) Lifetime distributions fit with a single exponential (noncognate, orange; diffuse-away green), or double exponential (cognate, blue) yielded one characteristic lifetime for non-cognate tethers (τ = 0.77 ± 0.05 s) and for diffusion away from *loxP* half-sites (τ = 1.83 ± 0.02 s). Two lifetimes are evident for particles that dissociate directly (τ_short_ = 2.0 ± 0.1 s and τ_long_ = 40 ± 24 s, 17 ± 3% population). (F) Probability of a particle adopting scanning and recognition states (*P_i_* = *A_i_τ_i_*/∑*A_i_τ_i_*) when localized at the *loxP* half site; particles located at the *loxP* half-site tend to be long-lived. (G) Fractional probabilities of direct loading at cognate sites versus one-dimensional diffusion from nonspecific DNA (N = 258). Uncertainties were estimated by bootstrapping with 1000 iterations.

Distinct residence times at cognate sites indicate Cre adopts both fast- and slow-scanning modes at these sites. To characterize recognition lifetimes, we constructed CDF plots for lifetimes in which Cre was located within ±0.1 μm of the center of the tether, corresponding to the location of the *loxP* half-site (**Figure 5E**). For tethers lacking the *loxP* half-site, we observed a single distribution with a characteristic lifetime of 0.77 ± 0.05 s, consistent with the time required to diffuse out of the 0.2 μm window. For tethers containing the *loxP* half-site, residence times were not well described by a single exponential. After segregating trajectories in which Cre was observed to slide away from the site from those in which it appeared to dissociate directly without sliding (**Figure S9**), distinct lifetimes were observed. Sliding away from the site could be described by a single exponential with a lifetime of 1.83 ± 0.02 s. The CDF of lifetimes for particles that dissociated directly are best described by a two-state distribution with characteristic lifetimes of 2.0 ± 0.1 s, and 40 ± 24 s, respectively, with the long-lived state corresponding to 17 ± 3 % of the traces (**Figure 5E-F**). Separate experiments showed that photobleaching is not significant at these timescales (Figure S6).

Bimodal lifetime distributions suggest that upon locating a cognate half-site, Cre may undergo a conformational switch between two recognition modes. To model the recognition process, we posited a two-step mechanism involving the sequestered and extended states, and assumed that direct dissociation from the extended conformation is negligible:

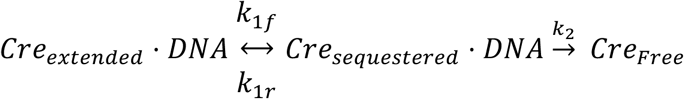

Here, *k_1f_* and *k_1r_* are the rate constants governing isomerization between the extended and sequestered conformations of Cre while bound to its cognate DNA, and *k*_2_ is the rate of dissociation from DNA of the sequestered state. We fit the experimentally determined 1-CDF lifetime distribution with an analytical expression derived for the case in which the starting point of the reaction is the DNA-bound sequestered conformation (included in the SI). This analysis yielded well-defined best-fit values for the rate constants: *k_1f_*: 0.03 ± 0.02 s^-1^, *k_1r_*: 0.08 ± 0.02 s^-1^ and *k_2_*: 0.43 ± 0.01 s^-1^. In this model, the ratio of the first two rate constants represents the equilibrium constant between the DNA-bound *extended* and *sequestered* conformations of Cre, revealing the extended conformation to be favored by *K =* 2.7 ± 1.8. This value may underestimate the true equilibrium since the 0.1 µM spatial resolution of our measurements is insufficient to distinguish particles that are bound at the cognate site from those that are proximal; the latter would inflate the population of short-lifetime localized particles. Nevertheless, the data are consistent with a model in which Cre locates *loxP* sites by adopting a sequestered, “scanning-competent” conformation and adopts an extended, immobile “recognition” conformation upon target site recognition.

Lastly, we examined whether Cre was more likely to bind DNA directly at the *loxP* half sites via direct three-dimensional diffusion or via one-dimensional scanning along the DNA. Direct loading was characterized by a trace being colocalized with the center of the tether in the first frame in which it was detected, while searching was characterized by first binding away from the *loxP* half site and diffusing to colocalize with the *loxP* half site. In these experiments, location of the half-sites was dominated by the one-dimensional “search” mechanism, with a probability of 0.7 ± 0.1 (**Figure 5G**).

This observation underscores the importance of efficient DNA scanning in site-selection by Cre.

## Discussion

### Insights into Cre-DNA Scanning from NMR and Single Molecule Tracking

We used single-molecule and ensemble methods to investigate a functional role for the cis-docked conformation in Cre recombinase. NMR and single-molecule tethered tracking experiments provide strong evidence that sequestration of the C-terminus serves to enable efficient sequence scanning by Cre along double-stranded DNA, while site recognition is accompanied by conformational changes that alter its diffusion properties. These observations reveal an unexpected function for the C-terminus of Cre, which has other essential roles in Cre-mediated recombination.

Heteronuclear ^15^N NMR relaxation experiments performed on the isolated Cre catalytic domain (**Figures 2, S2**) reveal that in the absence of DNA the enzyme undergoes reversible exchange between two conformations involving rearrangement of its C-terminal segment. Because the catalytic domain folds independently of the N-terminal domain (34), this two-state exchange behavior extends to full-length Cre. The major conformation in DNA-free Cre, favored by approximately 99% (∼5 *k_B_T*), corresponds to the sequestered *cis-*docked conformation previously identified through chemical shift and paramagnetic relaxation enhancement (PRE) NMR experiments. In this state, the C-terminal segment occludes the active site, thereby reducing the affinity of the protein for both cognate and non-cognate DNA (33). Exchange occurs with a transient minor conformation in which residues diagnostic for autoinhibition exhibit large differences in backbone amide ^15^N chemical shifts, consistent with a state where the C-terminus is in an extended-like conformation, as observed in DNA-bound structures. Thus, the data reveal a propensity in DNA-free Cre for a spontaneous conformational switch between a dominant sequestered conformation and a minor extended one.

Single-molecule fluorescence microscopy of Cre diffusing on DNA illuminated kinetic consequences of domain sequestration. Tethered tracking experiments showed that full-length Cre exhibits unconstrained random walk diffusion on double-stranded DNA, scanning approximately 2 kb per encounter (**Figure 3**). Conversely, deletion of the autoinhibitory sequence produces an enzyme that is essentially immobile and unable to scan along DNA to search for cognate DNA sites (**Figure 3, S5**). These observations indicate that domain sequestration enables dynamic DNA scanning, whereas its removal traps the enzyme on non-cognate sites, preventing 1D target search.

Solution NMR experiments of DNA-bound Cre provide insights into the conformation of diffusion-enabled Cre. Backbone amide NMR chemical shifts are highly sensitive to protein structure and environment. Upon binding to double-stranded DNA, chemical shift changes can be expected due to (i) direct changes in electronic environment for resonances in proximity to nucleic acid groups and (ii) indirect effects from conformational coupling to binding (43). By monitoring well-resolved signals diagnostic of the sequestered state (33, 34), we find that the full-length protein retains these diagnostic spectral signatures when bound to non-cognate DNA (**Figure 4**). In contrast, the spectral signatures disappear upon binding cognate DNA (**Figure S6**). These results suggest that efficient sequence scanning by Cre arises from its ability to preferentially populate the sequestered conformation only when bound to non-cognate DNA.

DNA target site recognition by Cre is accompanied by distinct changes in its diffusional behavior. Single-molecule tracking experiments using DNA tethers with a cognate *loxP* half-site at the center reveal two lifetimes: a short “scanning” mode and a long “recognition” mode (**Figure 5**). Upon encountering the *loxP* half-site, Cre is observed to pause at the site. When localized at the cognate site, Cre exhibits two distinct lifetime distributions, indicating dynamic exchange between the “recognition” and “scanning” states. We conclude that these distinct states reflect the role of domain sequestration in modulating the balance between rapid search and stable engagement at the target site.

### A Role for Domain Sequestration in DNA Site Selection by Cre Recombinase

In addition to playing a critical role in mediating cooperative DNA binding and synapsis by Cre, its C-terminal segment (helix αN in tetrameric structures (23)) appears to play an unexpected but critical role in mediating site-selection. Domain sequestration can be seen to act through both thermodynamic and kinetic effects (**Figure 6**). In the absence of DNA, ^15^N CEST NMR relaxation experiments reveal exchange between the autoinhibited major state and a minor extended conformation that is sampled transiently (**Figure 2**). The rate constant for association with DNA are estimated based on published kinetic analysis of time-resolved electrophoretic mobility shift experiments (44), and is assumed to be the same for cognate and non-cognate DNA. The rate constants for one-dimensional sliding on non-cognate DNA are derived from the fitted dissociation constants for WT and Δ300 Cre (**Figure 3**). We estimate upper limits for isomerization of Cre bound to non-cognate DNA from the observation of a single set of signals in NMR spectra, the signal-to-noise of the spectrum, and separation between corresponding signals assigned to the two states (**Figure 4**). Finally, ratios of the affinities of WT and Δ330 for a cognate half-site, suggest that the extended conformation is favored, while spectroscopic and structural data reveal a concomitant bend in the DNA (29, 33). In this model, the autoinhibited conformation (top) is favored on non-cognate DNA, exhibiting facile one-dimensional diffusion. Upon locating a *loxP* half-site, a conformational transition to the recognition mode (bottom) results in extension of the C-terminal segment and long lifetimes at cognate sites.

**Figure 6.**
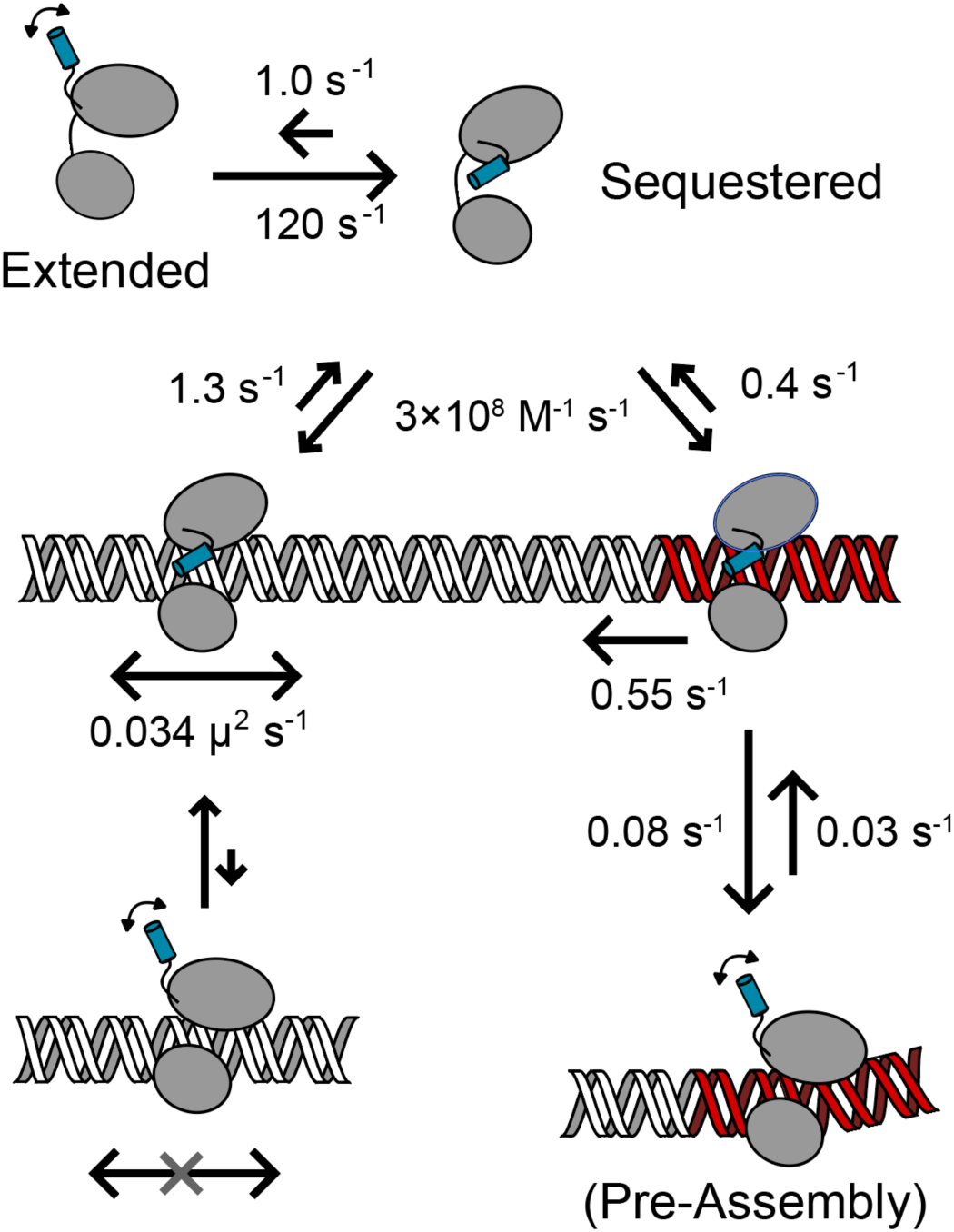
Model for the role of motif sequestration in site selection by Cre recombinase. Cre recombinase is shown schematically with core-binding and catalytic domains (ellipses) connected by a linker: the C-terminal tethering motif (blue cylinder) forms helix αN in tetrameric complexes. In the absence of DNA (top), Cre exchanges between sequestered (major) and extended (minor) conformations (*k*_ex_ ≈ 120 s^-1^, *p*_B_ < 0.01; Figure 2). The major sequestered state reversibly associates with DNA; rate constants for dissociation from non-cognate (grey) and cognate DNA (red) are derived from bound lifetimes in single particle tracking data (Figures 3, 5); the association rate is estimated from published results using mobility shift assays (44). The sequestered state scans DNA at rates obtained from the measured diffusion coefficients (Figure 3). At cognate sites, the extended conformation is favored, accompanied by modest DNA bending and arrested dissociation. At non-cognate sites, formation of the extended conformation would prevent scanning, and is strongly disfavored, consistent with NMR spectra (Figure 4). In this model, motif sequestration promotes diffusion on non-cognate DNA, while recognition of a *loxP* half-site triggers extension of the C-terminal segment and stable binding at cognate sites.

## Conclusion

Sequence-specific DNA-binding proteins face the fundamental challenge of rapidly locating rare target sites within vast genomes while avoiding stable engagement with the overwhelming excess of noncognate DNA. Here, we provide evidence that Cre recombinase addresses this challenge through a conformational equilibrium involving sequestration of its C-terminal tethering motif. NMR measurements reveal that, in the absence of DNA, Cre predominantly populates a sequestered state that transiently samples an extended conformation. Single-molecule tracking demonstrates that this autoinhibited state is not merely regulatory but is required for efficient one-dimensional diffusion along DNA. Removal of the C-terminal motif abolishes efficient scanning and traps the protein in an immobile DNA-bound state. These observations thus establish motif sequestration as a key determinant of DNA target search by Cre.

Our data further indicate that the conformational equilibrium governing DNA scanning is coupled to target recognition. NMR spectra show that the sequestered state persists when Cre interacts with noncognate DNA, whereas single-molecule measurements reveal prolonged residence times and distinct recognition states upon encounter with cognate *loxP* sequences. These findings are consistent with a model in which the sequestered conformation functions as a scanning-competent state that facilitates rapid exploration of DNA, while recognition of cognate sites is associated with a shift in the equilibrium toward an extended conformation that arrests diffusion and stabilizes site-specific binding. In this manner, a single conformational switch links rapid target search to selective target engagement.

Broadly, our results identify transient motif sequestration as a mechanism for balancing search efficiency with target specificity. Rather than serving as a passive constraint, the sequestered state actively promotes productive target localization while suppressing inappropriate stabilization on noncognate DNA. Similar search-versus-recognition transitions in other systems suggest that dynamic masking of DNA-binding surfaces may represent a general strategy for coordinating target search across diverse genomes. Ultimately, autoinhibition emerges not as a barrier to DNA recognition, but as the mechanism that enables efficient and selective target recognition.

## Methods

### Protein Purification

Plasmid containing wild-type (WT) Cre recombinase was generously provided by Gregory Van Duyne (University of Pennsylvania, Philadelphia, PA). All native cysteine residues were mutated using a Q5 site directed mutagenesis kit (New England Biolabs) with the following mutations: C49S, C55A, C155S, and C240A. For covalent labeling, a single cysteine was reintroduced using the mutation S226C. For Cre Δ330, a TAG stop codon was introduced following residue 330 as described previously (33) (E331STOP). Plasmids encoding the constructs were transformed into T7 Express-LysY/Iq *E. coli* cells (New England Biolabs) by electroporation and plated on LB media plates supplemented with 0.1 mg/mL carbenicillin for antibiotic selection. Single colonies from LB media plates were picked and used to inoculate 10 mL of LB media and 0.1 mg/mL carbenicillin and grown overnight shaking at 220 rpm at 37 °C. 10 mL of this culture was transferred to a 2 L flask containing 1 L LB media supplemented with 0.1 mg/mL carbenicillin for growth at 37°C shaking at 220 rpm. For [U-^15^N] Cre Δ330, 10 mL of the starter culture was added to 1 L of M9 minimal media supplemented with 1g ^15^NH_4_Cl (Cambridge Isotope Labs) and 0.1 mg/mL carbenicillin as described previously (33). Cultures were grown shaking at 220 rpm and 37 °C until the optical density at 600 nm reached 0.6. Once at sufficient optical density, 400 μL of 1 M IPTG (Isopropyl Beta-D-1-thiogalactopyranoside) was added and cells were allowed to induce for four hours while shaking at 220 rpm and 37 °C. Following the induction period, cells were pelleted by centrifugation at 4225 x g and stored at -80 °C. Protein was purified as described previously (33).

### DNA tether Purification

The Lambda DNA tether was prepared by filling the 12-base 5′ single-stranded overhangs on each end of the bacteriophage λ DNA with the DNA polymerase I Klenow fragment that lacks 5′→3′ endonuclease activity. The reaction solution contained the following reagents: 6 nM Lambda DNA, 100 µM dATP, 100 µM dTTP, 100 µM dGTP, 80 µM biotin-14-dCTP and 0.075 U/µL Klenow fragment in 1X NEB2 buffer. These reagents were incubated at 37 °C for 30 min followed by heat inactivation at 75°C for 20 min to complete the polymerase reaction. Using biotin-14-dCTP enables incorporation of 4-6 biotinylated dCTP during the polymerase reaction to provide the lambda tether with multiple biotins on each end. Klenow fragment was removed by phenol extraction followed by exchanging the tether into 0.5X TE (10 mM Tris pH 8.0 with 1 mM EDTA) at 4°C.

For *loxP*21L and NC-DNA custom inserts DNA tethers, insert sequences were designed to have 20 bp 5’-overhangs on each strand. Top and bottom strands were purchased from IDT and lyophilized ssDNA was resuspended in miliQ water. Top and bottom strands were mixed in equimolar ratios in buffer containing 10 mM TRIS pH 7.0 and 10 mM NaCl and were incubated in a water bath at 95 °C for 10 minutes. Following incubation, DNA was allowed to anneal by cooling slowly overnight. Annealed DNA was further purified by QHP chromatography (Cytiva) using a linear gradient (10 mM TRIS pH 7.0, 25 mM – 1 M NaCl). Purified DNA was visualized on a 2% agarose gel impregnated with 0.1% v/v Sybr Safe dye to assess purity (Invitrogen).

The 7.75 kbp DNA handle used in experiments involving the inserts were prepared by ligating biotin-modified and single stranded oligos to the ends of a linearized pPSU2 plasmid. Briefly, the plasmid was linearized using *Bsa*I to provide four-nucleotide non-palindromic sticky ends for ligation purposes. One end of the linearized plasmid was ligated to the following pre-annealed oligos to provide the biotin-modified end: 5′-BiotinTEG-AGCTAGCTTTCAATAGCTCG-3′ and 5′CGGTCGAGCTATTGAAAGCTAGCT-3′. Simultaneously, the other end of the linearized plasmid was ligated to the following pre-annealed oligos to provide the AGA CTC CGA CTG CCA GCC CA 5’-overhang: 5’-AGACTCCGACTGCCAGCCCACCTACACTGGGATAATTGAC-3’ and 5’-ACCGGTCAATTATCCCAGTGTAGG-3’. Following the ligation reaction, the enzymes were removed using phenol extraction, followed by exchanging the tether into 0.5X TE at 4 °C. Excess ligation oligos were purified away using anion exchange chromatography with a Gen-Pak^TM^ Fax 4.6 100 mm column (Waters) and buffer exchanged into the 0.5X TE for storage.

Biotinylated 7.75 kbp DNA handles with complimentary 5’-overhangs to the insert were mixed at a 6:1 handles:insert DNA ratio and incubated at 37 °C for 30 minutes in buffer containing 100 mM NaCl, 3.5 mM TRIS pH 8.0, and 350 μM EDTA. Annealing was confirmed by running a 0.7% agarose gel in TBE buffer. The assembled DNA complex was diluted to 0.1 nM in buffer containing 0.5X TE with 100 mM NaCl before loading onto the C-Trap.

NCD1 substrate for NMR was prepared as described previously (33).

### Optical Trapping and Single-Molecule Imaging

Single-molecule imaging experiments were performed on a Lumicks C-Trap instrument with a dual optical trap and microfluidics flow-cell. Cells were passivated with sequential washes of 1 mg/mL BSA and 0.5% (v/v) Pluronics F-127 solution (Invitrogen). Streptavidin coated polystyrene beads (2.14 and 4.34 μm diameters, Spherotech) were introduced into the system through channel 1 and captured using the 1022 nm IR-laser optical traps. Trap stiffness was calibrated in the absence of flow using a power spectrum and ranged from ∼0.2 – 0.4 pN/nm. DNA tether solution was added to channel 2. DNA tethers were captured between the beads and the complex was moved into channel 3 to ensure single DNA tethers were trapped by measuring force-extension curves resulting from repeated extension of the inter-bead distance and fitting to an extensible worm-like chain model (38) in the absence of flow. Cy5-labeled protein was diluted to 15-30 nM in imaging buffer (50 mM HEPES pH 7.4, 130 mM NaCl, 0.005% Tween-20, 2 mM Trolox, 450 µg/mL glucose oxidase, 22 μg/mL catalase, 0.0115% v/v COT (Cyclooctatetraen), 0.012% v/v NBA (3-Nitrobenzyl alcohol), 1.6% w/v glucose) and was introduced into channel 4 or 5 for observation. Using a 638 nm excitation laser at 10% power, two-dimensional scans were collected to set the confocal plane along the length of the DNA tether. Kymographs were acquired while holding the DNA at constant force of 10 pN, with a frame rate of 16-77 msec (longer framerates for longer λ DNA tethers). Kymographs were collected at multiple laser powers to ensure characteristic lifetimes were not limited by dye photobleaching.

### Data Visualization and Analysis

Kymographs were exported and visualized using Lumicks Lakeview Software.

Traces were tracked using the Python Pylake package (https://github.com/lumicks/pylake) and traces were exported for analysis using custom scripts. Briefly, the mean squared displacement (MSD) for each trace was measured using:

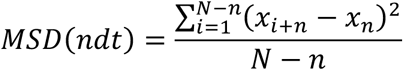

Diffusion coefficients and alpha values for each trace were determined using a double log plot:

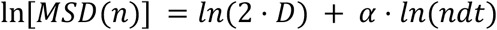

where n is an integer value for timepoints and dt is the frame rate of the kymograph. Points 3-23 (varying lag times between points) were used for fitting diffusive parameters as plots remain linear in this range (40, 57). Traces less than one second were omitted from analysis of diffusion coefficient and alpha values due to increased error in estimating diffusive values but were kept for fitting lifetimes. Alpha values were determined as the slope of the double log plot, while the y-intercept was used to calculate the diffusion coefficient at 1 sec. Lifetimes of traces were constructed using a cumulative distribution function (CDF) such that the maximum lifetime was set to zero and the median lifetime was set to 0.5. 1-CDF plots were fit using a single exponential decay function *A* ⋅ exp (−*τ*/*dwell*). The characteristic dwell-time was measured to be the reciprocal of the decay constant. Uncertainties in mean lifetimes were determined by bootstrapping 1000 times, retaining one-third of the data in each iteration.

Analysis of Cre bound to tethers containing either a *loxP* half site or noncognate DNA were visualized and traced using the same protocol as with lambda DNA. Traces were classified as interacting with the *loxP* half site if the center of the trace was within ± 0.1 μm of the center of the tether. To validate the presence of site-specific binding, the lifetime of all traces bound to the tether were fit to a single or double exponential decay, with uncertainties estimated by bootstrapping. To identify multistate binding to the *loxP* half site, lifetimes of traces only when bound to ± 0.1 μm of center of the tether were fit using a tripe exponential decay, with uncertainties estimated by bootstrapping.

### NMR Spectroscopy

Purified [U-^15^N]-Cre Δ330 was mixed with excess (2:1) noncognate DNA substrate, NCD1, at high ionic strength buffer (20 mM HEPES pH 7.0, 500 mM NaCl, 5 mM DTT). The complex was dialyzed stepwise into 1 L of buffer using 3.5 kDa MWCO dialysis cassettes (500 mM → 200 mM → 100 mM → 25 mM NaCl) to reduce the ionic strength and prevent aggregation of the complex. Once buffer exchanged, the complex was concentrated to a final volume of 250 μL and concentrations of 200 μM Cre + 400 μM NCD1 to ensure saturation (>99% at these concentrations). The sample was supplemented with 25 μL D_2_O (10% v/v) and 225 μM DSS (2,2-dimethyl-2-silapentane-5-sulfonate) for field locking and chemical shift referencing in a 3 mm NMR tube. ^1^H-^15^N TROSY-HSQC spectra were recorded using a Bruker Avance 800 MHz spectrometer with a triple-resonance inverse (TXI) cryoprobe. Spectra were recorded with 48 scans, 2048 x 256 complex points, and spectral widths of 21.85 x 36.00 ppm centered at 4.697 and 116 ppm for ^1^H and ^15^N respectively. Data was processed and visualized using NMRFx Analyst (58) (nmrfx.org).

### 15N Chemical Exchange Saturation Transfer

[U-^15^N]-CreCat (residues 127-343) was expressed and purified as described above for the full-length protein and was concentrated to 600 μM in NMR buffer containing 20 mM Hepes pH 7.0, 200 mM NaCl, and 5 mM DTT (dithiothreitol). The sample was supplemented with 10% D_2_0 (v/v) and 0.2 mM DSS for field locking and chemical shift referencing respectively. ^15^N amide chemical shift saturation transfer experiments were completed as described previously (37) on a Bruker Avance III HD 800 MHz spectrometer equipped with a 5 mm triple resonance cryoprobe with *z* axis gradients at 25 °C. A series of ^1^H -^15^N HSQC spectra were collected with ^1^H and ^15^N spectral widths of 16.02 x 28.0 ppm (12820.5 x 2270 Hz) for each offset frequency and an exchange time (T_Ex_) = 800 ms. For both B_1_ fields, 3 reference planes were collected with T_Ex_ = 0. This ensures the accuracy of the relaxation parameters derived from CEST(59). The B_1_ field calibration was carried out following previously established protocols (37, 60). This involved positioning the ^15^N carrier on an isolated resonance and incrementing T_rlx_ to acquire a free induction decay (FID). The oscillation frequency, determined via Fourier transformation or numerical fitting, was used to calculate the actual B_1_ field strength. For a B_1_ field of 22 Hz, 66 planes were collected with frequency offsets of 50 Hz (0.62 ppm), 8 scans per plane, and 2048 x 192 points. For a B_1_ field of 44 Hz, 33 planes were collected with frequency offsets of 100 Hz (1.23 ppm), 32 scans, and 2048 x 100 points in the direct and indirect dimension (61). The resulting pseudo-3D spectra were processed using NMRPipe (62) and the ratio of peak intensities were measured using the Autofit.tcl script. CEST profiles were fit using ChemEx (https://gbouvignies.github.io/ChemEx/) to a two-state exchange model. During global fitting, two rounds of fitting were employed. In the first round *k*_ex_, *P*_B_, and δω_AB_ were held constant while the chemical shift of the minor state (δ_B_), *R*_1A_, and *R*_2A_ were fit. During the second round, the previously fit values were held constant while k_ex_, P_B_ and δω_AB_ were fit. Uncertainties were estimated using Monte Carlo simulations based on signal-to-noise ratios.

## Acknowledgements

This work was supported by NIH R01 GM122432 (to M.P.F) and NIH R35 GM139564 (to M.G.P.). J.S.M was supported by NIH T32 GM144293. Analysis of NMR data was done using NMRbox: National Center for Biomolecular NMR Data Processing and Analysis, a Biomedical Technology Research Resource (BTRR), which is supported by NIH grant P41GM111135 (NIGMS). Support and instrumentation for the LUMICKS C-TRAP was made possible by NIH S10 OD028705. Partial funding for the OSU NanoSystems Laboratory shared facilities used in this research was provided by the Center for Emergent Materials: an NSF MRSEC under award number DMR-2011876.

