## Supporting Information for "Enhanced DNA Scanning and Target Specificity via Motif Sequestration"

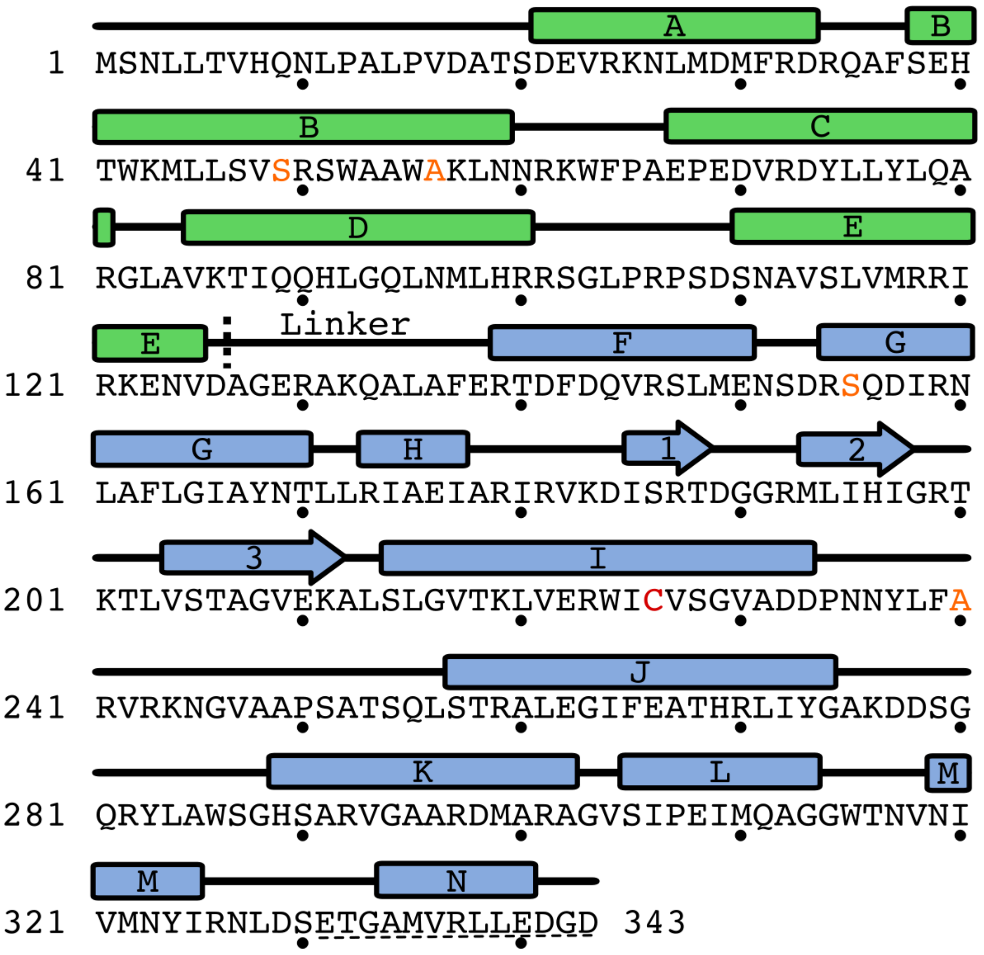


**Supplemental Figure 1. Sequence map of Cre recombinase and mutations introduced in the current study.** Secondary structure map of the core binding (residues 1-126) and catalytic domains (Cre$^{Cat}$, 127-343) are shown in green and blue respectively. Dashed line shows beak between two domains. Residues 330-343 constitute the inter-protomer tethering motif. Positions of native cysteines are denoted in orange as the new amino acid identity. Position of the S226C mutant used for Cy5 labeling is highlighted in red. The sequence deleted in the Δ330 mutant is underlined.

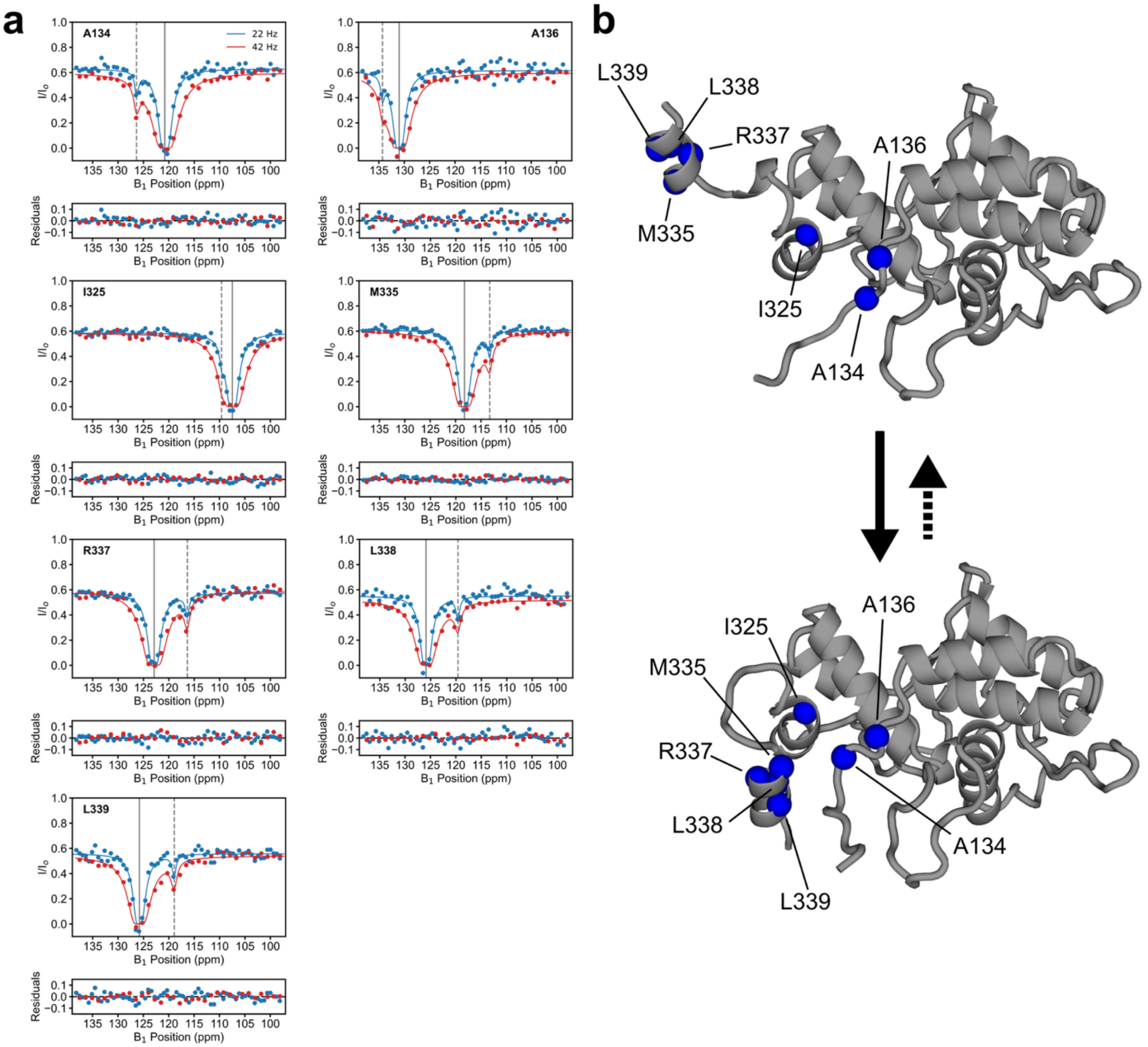


**Supplemental Figure 2. Backbone amide $^{15}$N resonances of many residues in Cre exhibit NMR CEST profiles indicative of slow two-state exchange**. (a) All CEST profiles in Cre$^{Cat}$ that exhibit two-state exchange. Data were globally fit with the same two-state model to extract the exchange rate and populations. (b) Model for the exchange probed in the CEST experiments. (Top) isolated catalytic domain from 2HOI chain A and (bottom) NMR PRE derived model (1) for the sequestered conformation of the catalytic domain.

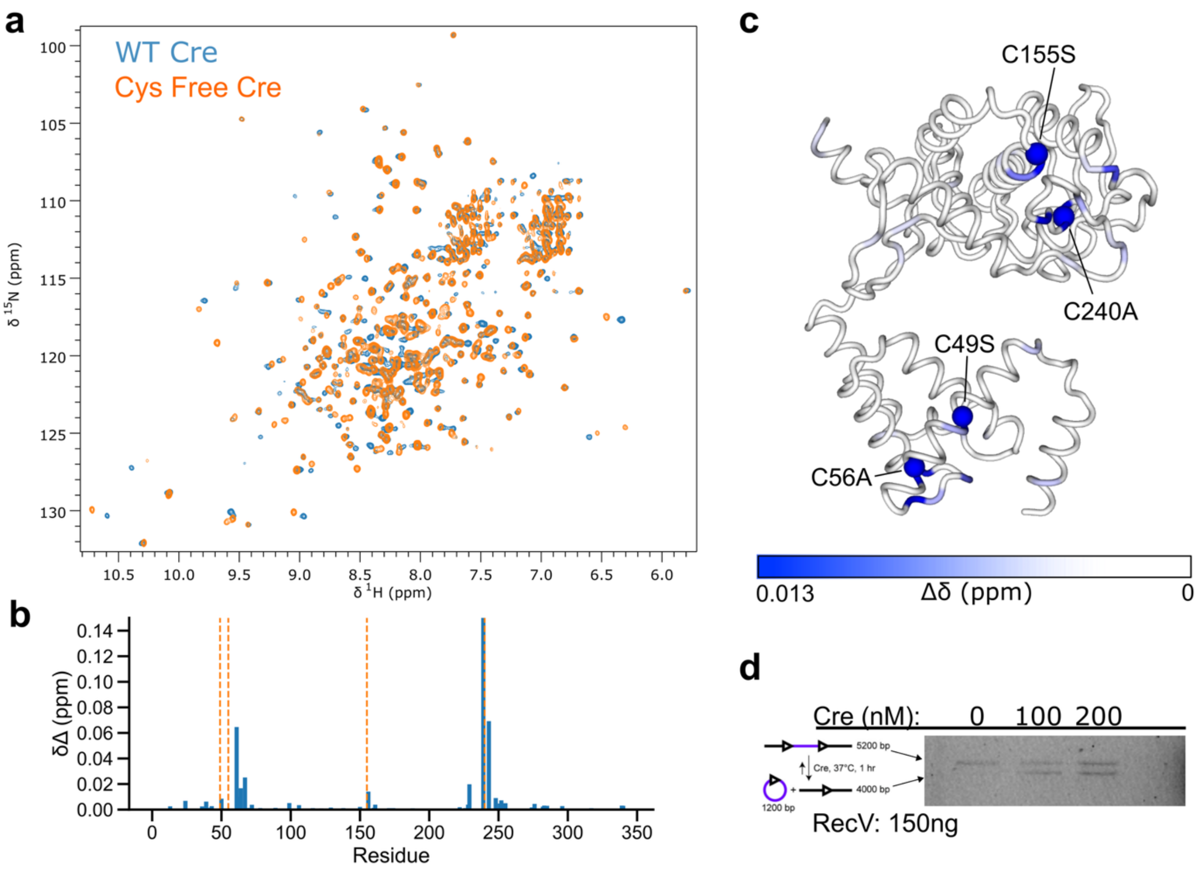


**Supplemental Figure 3. Replacement of the four native cysteine residues in Cre does not significantly alter its structure or impair activity**. (a) TROSY HSQC Spectra of full-length WT Cre (blue) overlayed with cysteine-free Cre. (b) $^{1}$H-$^{15}$N Chemical Shift perturbations for assigned to residues in Cre. Orange dashed lines indicate the location of the cysteine mutations. (c) Combined amide $^{1}$H-$^{15}$N chemical shift perturbations Δδ mapped onto the structure of Cre (PDB 2HOI, chain A), using $\Delta\delta = \sqrt{\Delta\delta_H^2 + \Delta\delta_N^2/25}$. Chemical shift perturbations are mostly located at the sites of mutations, indicating not global effect on structure. (d) *In vitro* recombination activity of cysteine-free Cre was assayed using 150 ng of linearized plasmid containing two *loxP* sites separated by 1200 bp.

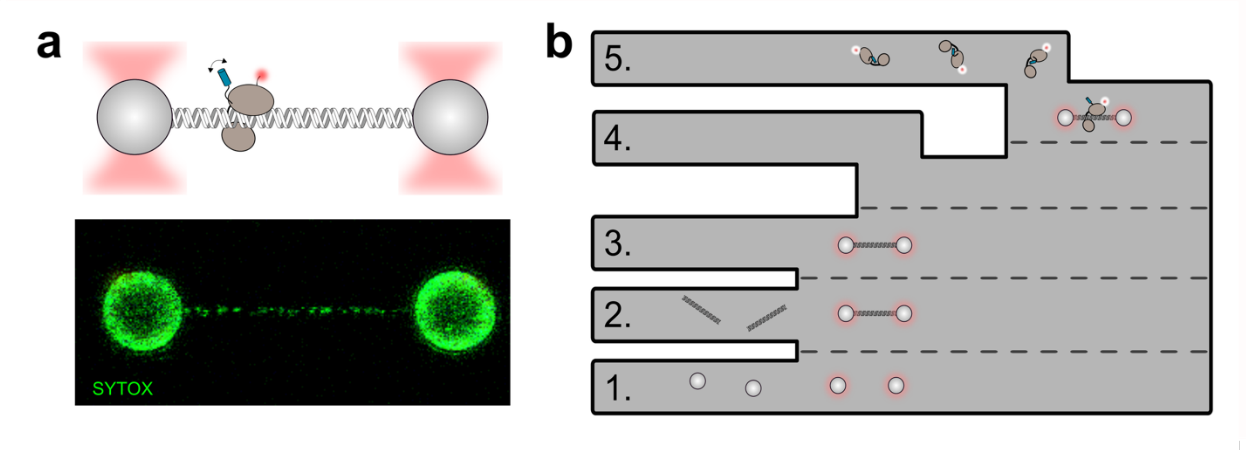


**Supplemental Figure 4. Fluorescence microscopy and optical tweezers setup**. (a) Illustration of experimental setup in which streptavidin coated polystyrene beads are captured in a dual optical trap, and a biotinylated lambda DNA tether is bound between the beads. Fluorescently labeled Cre can then be observed on the DNA tether. Using the fluorescent intercalating agent Sytox, we can directly observe the DNA tether between the beads. (b) Diagram of the microfluidics cell: polystyrene beads are first captured in channel **1** using the optical traps before being calibrated in channel **3**. Beads were then moved to channel **2** to capture a DNA tether and moved to channel **3** to ensure only a single tether was present. Once the ternary bead-tether complex was formed, it is moved to channel **5** in which fluorescently labeled Cre is introduced for observation. Experiments were conducted using a LUMICKS, Inc C-Trap instrument.

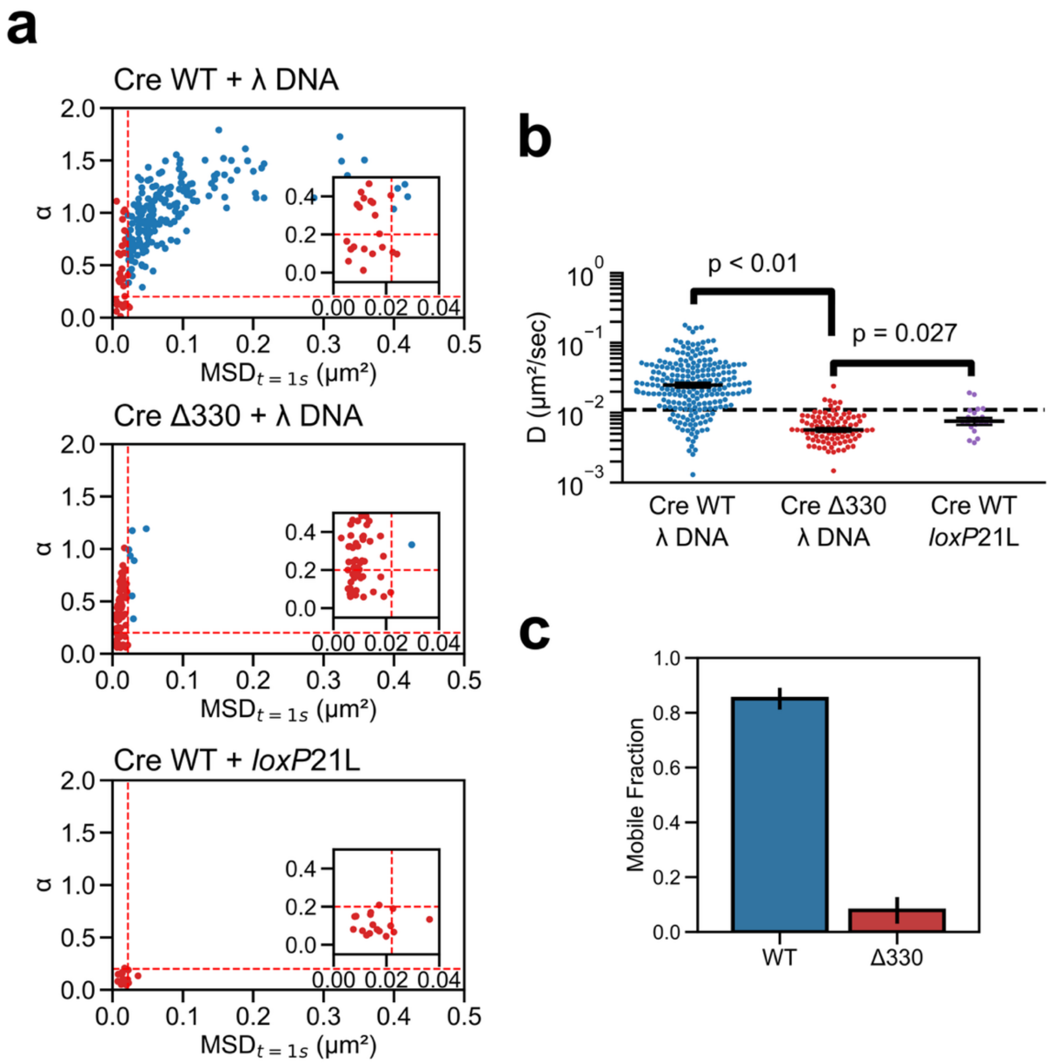


**Supplemental Figure 5. Cre Δ330 is predominantly immobile on lambda DNA.** (a) α versus MSD Correlation plots for WT Cre on λ DNA (top), Cre Δ330 on λ DNA (middle), and Cre on a tether containing a *loxP* half-site (bottom). Dashed red lines indicate cutoff values for non-diffusive particles, as determined for immobile nucleosomes (2). Datapoints in red indicate α and diffusion coefficients below the threshold. Mobility of particles bound to the cognate half-site was assessed for traces in which particles colocalized with the *loxP* half site location for more than 20 seconds. (b) MSD values at one second time lag for WT Cre on λ DNA (blue), Cre Δ330 on λ DNA (red), and WT Cre colocalized with the cognate half site for more than 20 seconds. (c) Percentage of traces classified as mobile as determined by both the α and *D* being above the threshold values.

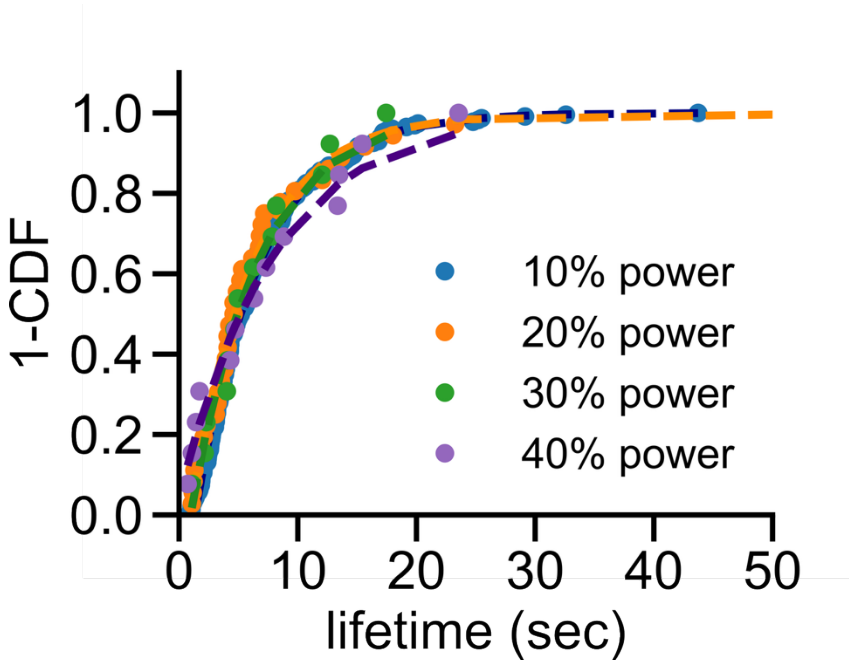


**Supplemental Figure 6. Lifetimes on DNA are not photobleaching limited.** To assess whether observed lifetimes on lambda DNA are due to dissociation or dye photobleaching, kymographs were collected at a range of excitation powers. We observed that laser power does not influence that overall lifetime distribution, indicating that the observed lifetimes are reflective of bound lifetimes on lambda DNA.

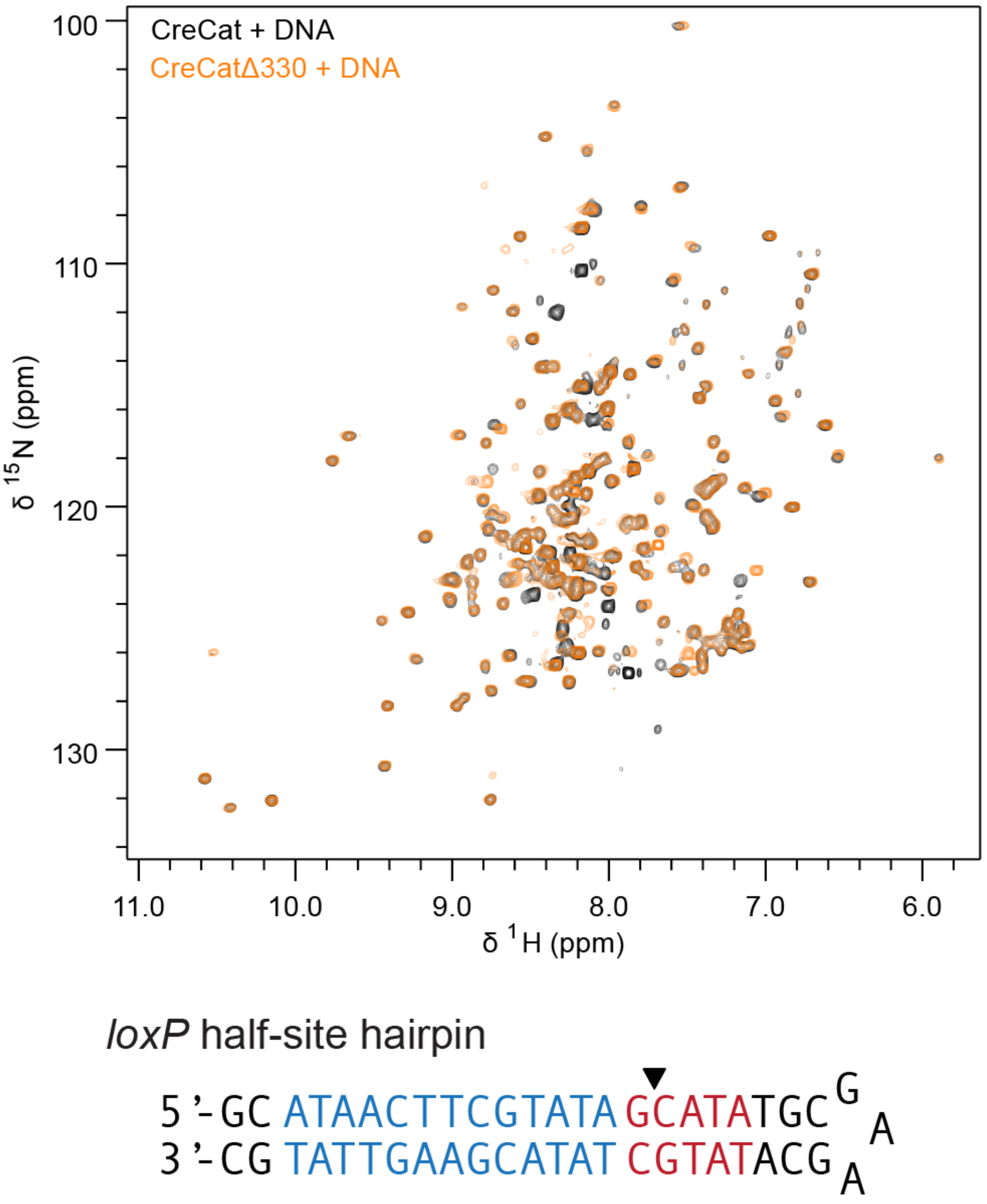


**Supplemental Figure 7. Cre and Δ300 adopt the same recognition structure on a *loxP* DNA half-site.** Top: Overlayed TROSY-HSQC spectra of [U-$^{15}$N] WT Cre$^{Cat}$ (residues 127-343, black) and [U-$^{15}$N] Cre$^{Cat}$-Δ330 (residues 127-330, orange) in the presence of a saturating concentration of a *loxP* half-site DNA hairpin substrate (Bottom). NMR spectra were recorded at 800 MHz, 25°C, in buffer containing 10 mM Tris, 100 mM NaCl, and 0.02% $NaN_3$, pH 7.0. (1)

a

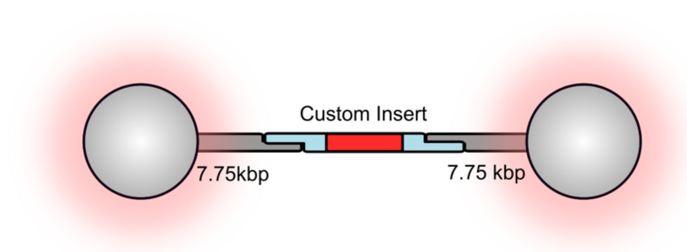


b

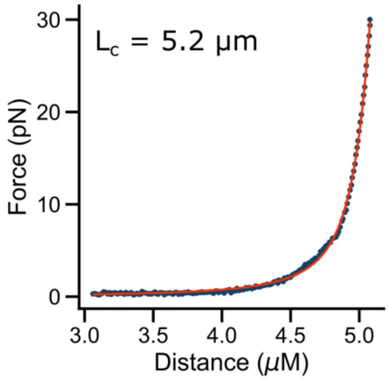


c

**Handle 5' overhang:**

```
5'- AGA CTC CGA CTG CCA GCC CA -3'
```

**loxP 21L half site:**

```
TS: 5' - TGGGCTGGCAGTCGGAGTCTGCATGCTAACCATAACTTCGTATAATGTCGTACGTAACCG - 3'
BS:                         3' - CGTACGATTGGTATTGAAGCATATTACAGCATGCATTGGCTCTGAGGCTGACGGTCGGGT - 5'
```

Noncognte (NCD1):

```
TS: 5' - TGGGCTGGCAGTCGGAGTCTGCATGCTAAGCGATCGAGTACTGCTAGCGGTACGTAACCG - 3'
BS:                         3' - CGTACGATTCGCTAGCTCATGACGATCGCCATGCATTGGCTCTGAGGCTGACGGTCGGGT - 5'
```

**Supplemental Figure 8. Design of DNA tethers containing custom DNA insert segments.** (a) Schematic of custom tether design. 7.75 kbp biotinylated DNA handles are attached to 2.14 µm streptavidin coated beads. Handles anneal to a synthetic DNA insert with single-strand DNA regions that share homology to the 5' overhangs on the DNA handles. (b) Force extension curve indicates single tether binding with the expected contour length. (c) DNA sequences for the 5' overhang, loxP21L (*loxP* half site) insert, and NCD1 (noncognate) insert. For the *loxP*21L insert, the RBE is shown in blue and the truncated spacer sequences is shown in red. 5' overhangs for the inserts are shown in orange.

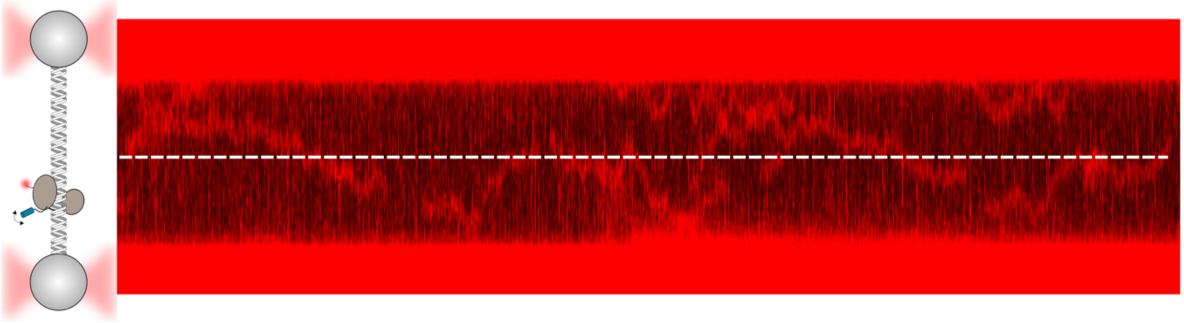

**Supplemental Figure 9. Examples of Cre crossing over the noncognate DNA insert**. Representative traces of Cre bound to the noncognate DNA insert tether illustrating crossover and transient interactions with the insert sequence consistent with a bypassing state. White dashed line indicates the position of the insert containing noncognate DNA.

### Analytical solution to consecutive reaction with an intermediate

We considered consecutive two-step reaction with a reversible first step:

$$A \underset{k1r}{\overset{k1f}{\rightleftharpoons}} B \overset{k_2}{\rightarrow} C$$

This can be modeled using a system of coupled linear first-order differential equations:

$$\frac{d[A]}{dt} = -k_{1f}[A] + k_{1r}[B]$$

$$\frac{d[B]}{dt} = k_{1f}[A] - (k_{1r} + k_2)[B]$$

$$\frac{d[C]}{dt} = k_2[B]$$

The equations for [A] and [B] can be written in matrix form:

$$\frac{d}{dt}\begin{bmatrix}[A]\\ [B]\end{bmatrix} = \begin{bmatrix}-k_{1f} & k_{1r}\\ k_{1f} & -(k_{1r}+k_2)\end{bmatrix}\begin{bmatrix}[A]\\ [B]\end{bmatrix}$$

The integrated solution is a linear combination of its eigenvalues, the decay constants $\lambda_1$ and $\lambda_2$, and their corresponding pre-exponential eigenvectors ($\vec{v}_1, \vec{v}_2$):

$$\begin{bmatrix}A(t)\\ B(t)\end{bmatrix} = c_1\vec{v}_1 e^{\lambda_1 t} + c_2\vec{v}_2 e^{\lambda_2 t}$$

The decay constants, $\lambda_1$ and $\lambda_2$, are the roots of the characteristic quadratic equation derived from the system's matrix:

$$\lambda^2 + (k_{1f} + k_{1r} + k_2)\lambda + k_{1f}k_2 = 0$$

The two distinct, negative roots of the quadratic are:

$$\lambda_1 = \frac{-(k_{1f} + k_{1r} + k_2) + \sqrt{(k_{1f} + k_{1r} + k_2)^2 - 4k_{1f}k_2}}{2}$$

$$\lambda_2 = \frac{-(k_{1f} + k_{1r} + k_2) - \sqrt{(k_{1f} + k_{1r} + k_2)^2 - 4k_{1f}k_2}}{2}$$

Applying t=0 boundary conditions: $[B]_0 = B_0$ and $[A]_0 = [C]_0 = 0$ yields the following analytical expressions for the time-dependent concentrations of *A* and *B*:

$$[B](t) = \frac{B_0}{\lambda_1 - \lambda_2}\left[(\lambda_1 + k_{1f})e^{\lambda_1 t} - (\lambda_2 + k_{1f})e^{\lambda_2 t}\right]$$

$$[A](t) = \frac{k_{1r}B_0}{\lambda_1 - \lambda_2}\left(e^{\lambda_1 t} - e^{\lambda_2 t}\right)$$

Then, using mass balance $[C](t) = B_0 - [A](t) - [B](t)$, and simplifying, yields:

$$[C](t) = B_0\left[1 + \frac{k_2}{\lambda_1 - \lambda_2}\left(\left(1 + \frac{\lambda_1}{k_{1f}}\right)e^{\lambda_1 t} - \left(1 + \frac{\lambda_2}{k_{1f}}\right)e^{\lambda_2 t}\right)\right]$$

This function was coded in a Python Jupyter notebook in Google's Colab environment, assisted by the Gemini agent, and the top 95% of the data were fit using the SciPy curve_fit module. Parameter uncertainties were computed from the resulting covariance matrix, yielding values of $k_{1f}$ = *0.03* ± 0.02 $s^{-1}$; $k_{1r}$ = 0.08 ± 0.02 $s^{-1}$; $k_2$ = 0.43 ± 0.01.

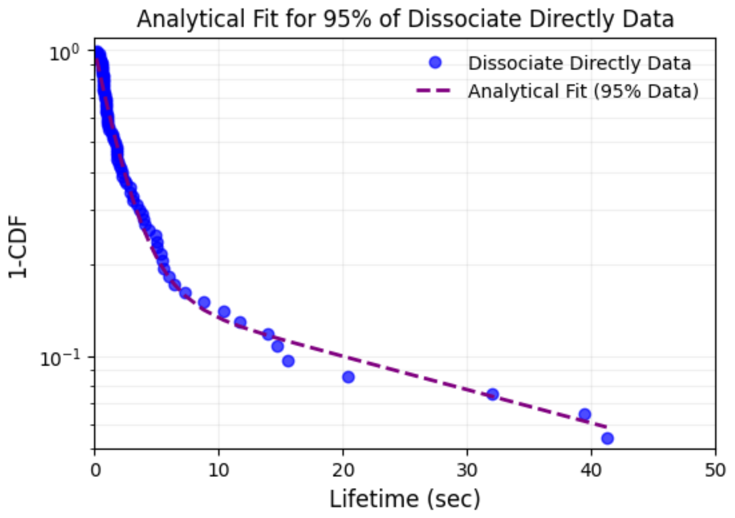
